# Syrah, a Slide-seqV2 pipeline augmentation

**DOI:** 10.1101/2022.03.20.485023

**Authors:** C.E. Brewster, F.G. Mann, B.W. Benham-Pyle, A. Sánchez Alvarado

## Abstract

Spatial transcriptomic techniques such as Slide-seqV2 uncover novel relationships and interactions between cell types by coupling gene expression and spatial data. Here we discuss two unexpected sources of error in Slide-seqV2 data, one physical and one computational. To address this we present an analysis pipeline augmentation, Syrah, which corrects for these errors and show that it improves both data quantity and quality over the standard pipeline alone or in combination with additional sequencing.

## Introduction

By capturing gene expression data coupled to spatial information, spatial transcriptomics provides novel biological insights about tissue composition and cell-type interactions. Slide-seqV2 is a spatial transcriptomic method that uses uniquely barcoded oligonucleotide covered beads arranged in a monolayer on a glass slide to capture mRNAs from their immediate vicinity [6]. Because the positions of the beads on the slide have already been established, bead identity can be used to assign spatial coordinates to each read. Positional and molecular identifiers are incorporated from the bead during reverse transcription and paired-end sequencing determines the positional and molecular identifiers in read 1 and mRNA-sequence data in read 2.

Unbiased spatial transcriptomics methods present unique opportunities for scientists working on research organisms where genetic markers and tissue architectures are less characterized. We used Slide-seqV2 to generate a spatial transcriptomic atlas of the free-living freshwater planarian, *Schmidtea mediterranea*, a model for tissue regeneration [4]. Regenerating tissue fragments were arranged around an intact animal, embedded in histology media, and a single cryo-section was used for Slide-seqV2. During our analysis, we discovered four types of errors: lost reads, lost beads, duplicated reads, and reads assigned to the wrong bead. We identified two likely sources of these errors, one arising from the synthesis efficiency of the mRNA capture oligonucleotides [1, 2], and the other a computational artifact of determining bead position using SOLiD sequencing [3]. We created an analysis pipeline augmentation (Syrah) to correct for these errors, which significantly improved the quantity and quality of data.

### Oligonucleotide synthesis errors cause frameshifts in bead barcodes

Slide-seqV2 captures mRNA using a bed of beads coated in oligonucleotides that are synthesized *in situ* on the bead surface via reverse-direction phosphoramidite synthesis, which adds one nucleotide per cycle from the 5’ to the 3’ end [5, 6]. During each cycle of synthesis there is a chance that the intended nucleotide will fail to be incorporated [1, 2], and when this occurs it causes a “deletion” in the final oligonucleotide [Fig 1A]. As a result, the remaining sequence after the deletion is shifted toward the 5’ end.

**Figure 1:**
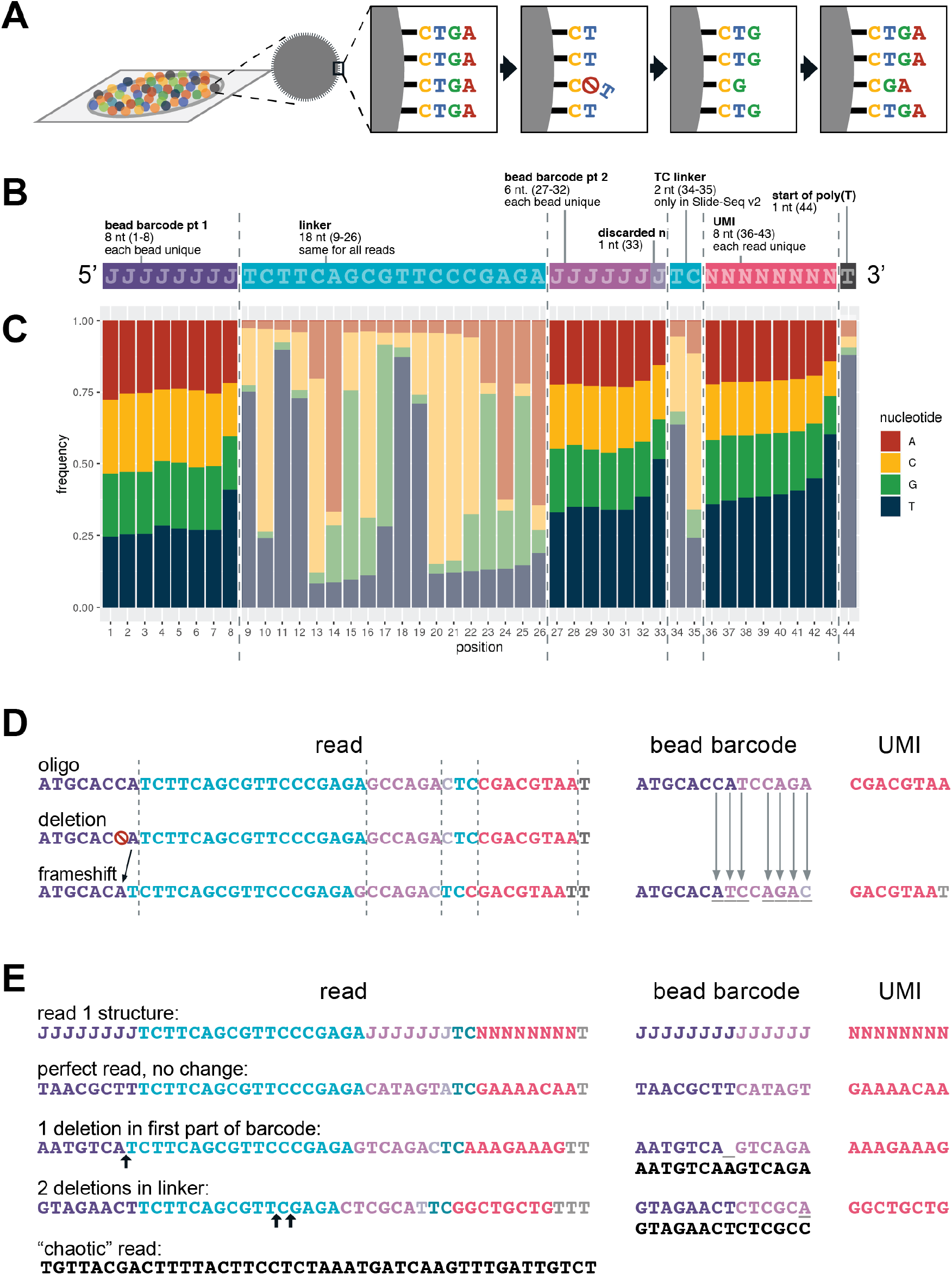
(A) Slide-seqV2 oligonucleotide synthesis, showing deletions caused by failed nucleotide incorporation. (B) Structure of the oligonucleotide that forms read 1 in Slide-seqV2 [Supplementary section 1]. (C) Nucleotide frequencies for the first 43 positions in a random 1 million read sample of the read 1 data showing evidence of increasing thymine bias toward the 3’ end. (D) A single deletion in the first barcode region of an oligonucleotide results in a frameshift that causes incorrect regions of the sequence to be interpreted as the bead barcode and UMI, resulting in a mismatch between the extracted bead barcode and the actual bead barcode. (E) The read 1 structure (top) compared with the Syrah’s output for reads of different types.

Each oligonucleotide on a Slide-seqV2 bead puck contains three types of sections: 1) the unique molecular identifier (UMI) which should be specific for each molecule on a bead; 2) the barcode (in two parts) which should be the same for all molecules on a given bead; and 3) the invariant sections – such as the linker – that should be the same for all molecules on the puck [Fig 1B]. Each oligo ends in a poly(T) tail, so deletions cause an increasing thymine bias toward the 3’ end. Since the first 44 positions of this oligonucleotide are sequenced for read 1, we can observe this 5’ thymine bias by looking at relative abundances of the four nucleotides at each position [Fig 1C]. The overabundance of thymine at position 43 (*T*_43_) can then be used to estimate the rate of deletions (*R_del_*) in the oligos using the formula [Supplementary section 1]

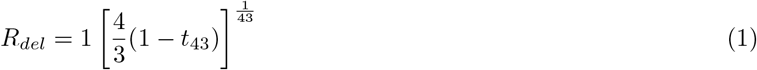

When applied to our dataset, the estimated deletion rate was approximately 1.47%.

The standard Slide-seq analysis pipeline extracts the bead barcode and UMI from the expected positions [6], and in the case of reads with deletions this results in the wrong barcode sections being extracted [Fig 1D]. Extracted barcodes are compared against the barcodes on the puck by computing the Hamming distance between the two barcodes: the number of positions at which the barcodes differ. Because deletions effectively cause a frameshift, a single deletion early in the barcode can cause many mismatches [Fig 1D]. If the barcode read does not match a barcode on the puck, the standard pipeline discards the data. On the other hand, if the incorrectly extracted read does match a barcode on the puck, the data is assigned to the wrong bead. In this dataset, for example, the barcode TTTTTTTTTTTTTT has over 13,000 reads assigned to it, while the next highest has less than half of that, strongly indicating that reads can be falsely assigned to beads.

To correct for synthesis-derived deletions in read 1 sequences, Syrah aligns the known linker sequence to each read and uses the beginning and end points of that alignment to determine where to extract the barcode and UMI segments. If the linker aligns perfectly, the read is processed the same as in the standard Slide-seqV2 pipeline. If there is a deletion in the first half of the barcode, a shortened barcode part 1 is extracted, and the barcode part 2 and UMI are extracted from one position to the left. If there are one or more deletions in the linker, the positions for the barcode part 2 and UMI are changed accordingly. Only when the linker cannot be satisfactorily aligned to a read is that read is discarded. Finally, when read barcodes are matched to puck barcodes, the distance metric allows for both substitutions and deletions [Fig 1E] in order to increase accurate matching between the sequenced read and the predicted bead barcodes.

In our data, roughly 56% of reads are “perfect” reads (no deletions in barcode part 1 or linker) which required no correction, 18% of reads have one deletion in the first part of the barcode, 13% of reads have between one and eight deletions in the linker (a read may have a deletion in the first part of the barcode and in the linker), and 17% of reads are “chaotic” reads with no acceptable linker alignment [Table S1].

### Positional sequencing ambiguity leads to bead “forking”

After Slide-seqV2 pucks are made, the position of each bead is determined using non-destructive in situ sequencing by oligonucleotide ligation and detection (SOLiD) sequencing [3, 5]. When the image analysis pipeline cannot differentiate between two possible nucleotides at a given position, a new virtual “bead” is computationally created with nearly identical x,y coordinates to the original. Then one of the two possible nucleotides is assigned to the original bead, the other nucleotide is assigned to the newly-created “bead”, and sequencing continues. This “bead forking” can occur more than once for each original (physically extant) bead.

Bead forking can lead to data loss. During analysis, reads may match and be assigned to either of these bead barcodes, with observed proportions ranging from nearly 50/50 to 5/95 or more. If reads are split between forked beads such that neither has enough reads to pass the data quality threshold, all reads from the original bead are lost. If only one bead passes the quality threshold, some data is lost. However if both beads pass the quality threshold, each bead will have its UMIs de-duplicated separately, potentially resulting in duplicated gene features.

Syrah corrects this via “de-forking.” Forked beads are identified by their physically impossible x,y proximity and barcode matching distance. Then reads assigned to any of those barcodes are redirected to a single barcode in the fork chosen arbitrarily.

In this data 37.5% of all bead barcodes in the puck coordinate file are part of a fork and de-forking resulted in significantly fewer bead overlaps in the predicted bead distribution [Fig 2E]. Comparison across multiple pucks indicates that estimated oligonucleotide deletion rate and bead forking rate may be correlated (data not shown).

**Figure 2:**
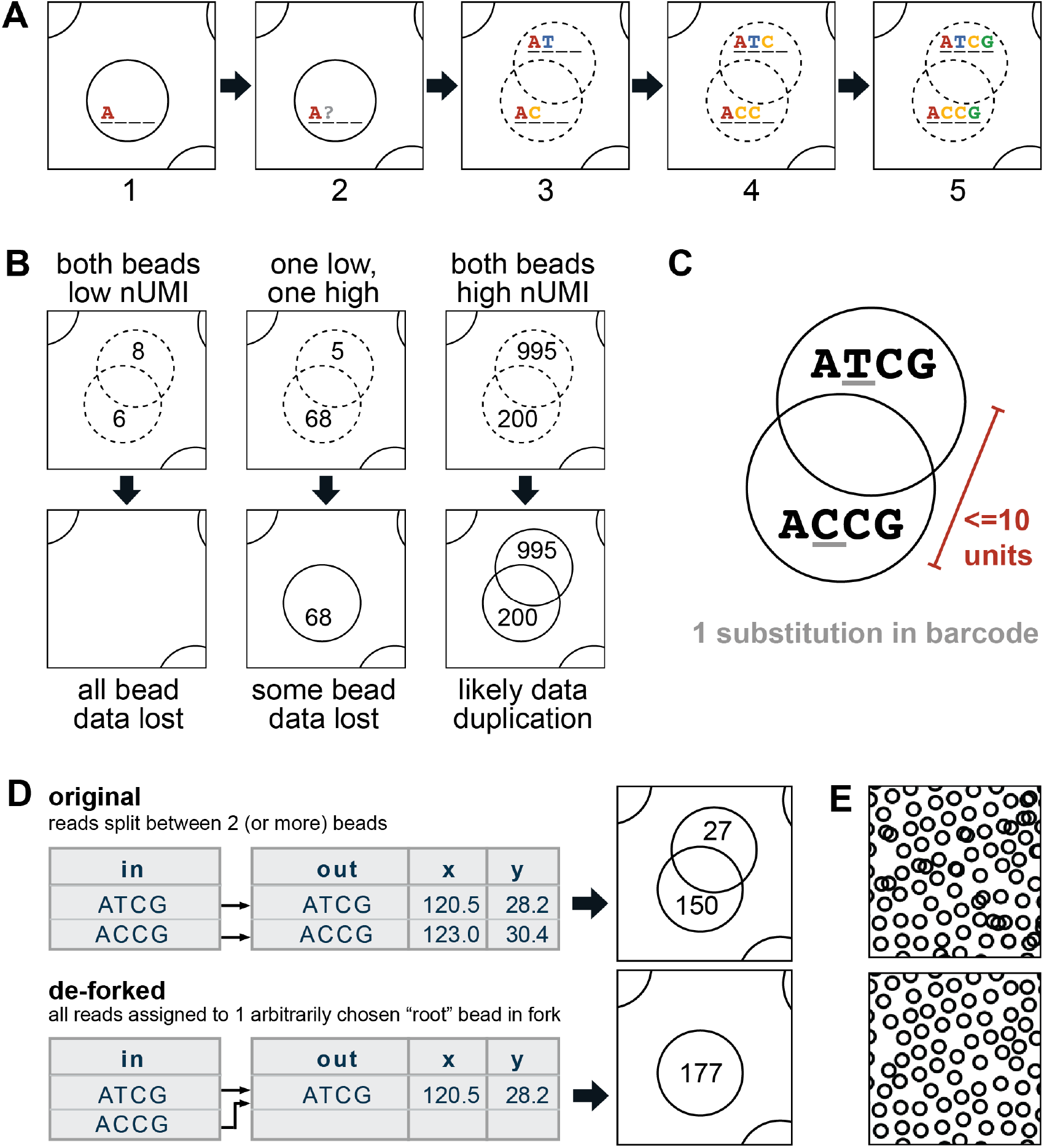
(A) Bead position on the puck is determined one nucleotide at a time via non-destructive sequencing. When a position on a bead cannot be determined with high confidence, the bead is split into two beads with similar positions, each of which is assigned one of the two possible nucleotide identities for that position, and sequencing continues. (B) Forked beads can result in several kinds of errors depending on the number of UMIs and their distribution across beads in the fork. (C) Forked beads are defined as being within 10 x,y units of each other on the puck and having barcodes differing at only one position. (D) Reads from forked beads are split between beads during barcode mapping with the standard pipeline (top) but are all assigned to a single “root” bead in the fork by Syrah (bottom). (E) Representative bead distribution on the puck, showing a 10 unit diameter, which is the expected size of the bead. Top is original data, bottom shows the same region after de-forking.

**Figure 3:**
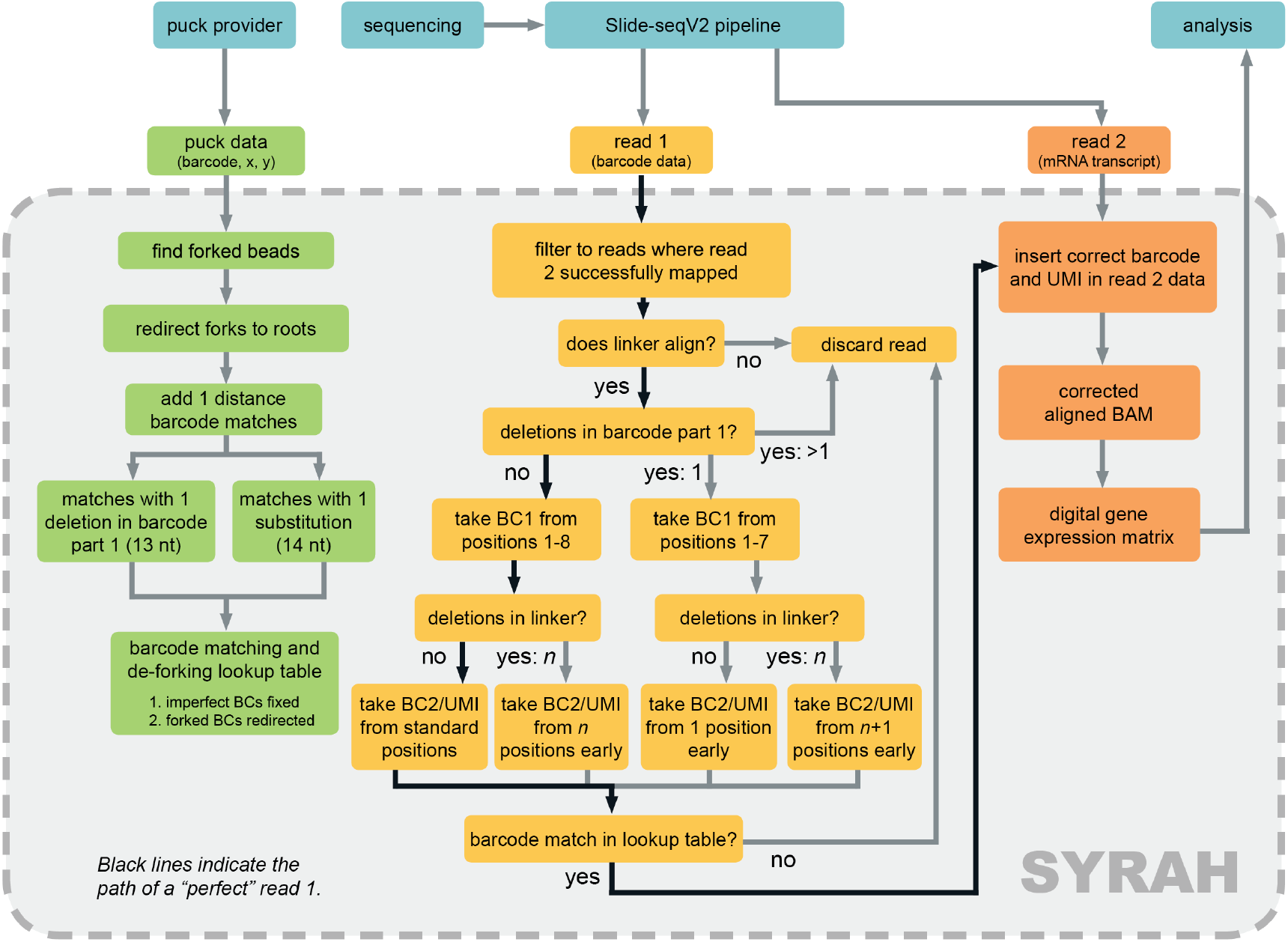
Schematic of Syrah’s read processing and its integration into the standard Slide-seqV2 workflow.

### Syrah corrects for read 1 deletions and bead forking

Syrah was built as an augmentation to the original Slide-seqV2 pipeline, such that it takes as input the output from the original pipeline and creates a corrected version of the data, facilitating comparison with the original pipeline’s results.

The first step of Syrah builds the barcode matching table that will be used to de-fork beads and perform bead barcode matching in a single step. Building this table requires only the puck coordinate file provided with the puck and can be performed prior to receiving sequencing data. Forked beads are identified as being within 10 units of each other on the slide and have barcodes that are a single substitution apart. Both barcodes are set to match to a single one of the two during barcode matching. Because the x,y coordinates are so similar, choosing an arbitrary member of the fork to assign the reads to does not meaningfully affect downstream analysis. This process is repeated iteratively to resolve multi-forked beads. Next, the possible one-distance matches are created, by generating all unique matches with one substitution anywhere in the barcode (14 nucleotide matches) or one deletion in the first half of the barcode (13 nucleotide matches). When read 1 barcodes are matched, they are error-corrected and then the corrected barcodes are de-forked.

Syrah’s second step corrects the extracted bead barcode and UMI in the aligned data. This requires both read 1 and reference-aligned read 2 data. For each pair of reads, the known linker sequence is aligned to read 1, allowing for up to one deletion in the first part of the bead barcode, and a total linker alignment distance of up to five [Supplementary section 3], including no more than one substitution. Reads without acceptable linker alignments are discarded. The linker alignment start and end positions are used to determine the positions from which the barcode and UMI are extracted. If the barcode matches successfully, the new barcode and UMI are substituted into the reference-aligned read 2 data, which is then written out to a corrected file.

After this, headers from the original aligned read 2 BAM file are added back in and the resulting SAM file can be converted to a digital gene expression matrix for downstream analysis.

### Evaluating Syrah

We evaluated the effects of Syrah relative to additional sequencing using Slide-seqV2 data generated from regenerating *S. mediterranea* animals. Four versions of the data are compared here: 1) one sequencing run of data processed using the standard pipeline; 2) two sequencing runs of data processed using the standard pipeline; 3) one sequencing run of data processed with Syrah, and 4) two sequencing runs of data processed with Syrah.

### Data generation

Animal husbandry, tissue handling and sectioning, puck synthesis, library preparation and sequencing were all performed per Benham-Pyle et al. (IN PREPARATION).

Following sequencing, Illumina Primary Analysis version NextSeq RTA 2.4.11 and Secondary Analysis version bcl2fastq2 v2.20 were run to demultiplex reads and generate FASTQ files, which were then run through the standard Slide-seq pipeline [5] using default settings. Code was retrieved from the Macosko Lab GitHub page: https://github.com/MacoskoLab/slideseq-tools. Reads were mapped to the Sánchez Alvarado lab transcriptome [7], both mapped read 2 data and unmapped read 1 data were processed using the pipeline modification with a maximum of one deletion in the barcode, maximum five linker alignment distance, and maximum one barcode matching distance. For all datasets only beads with at least 30 UMIs were included.

## Results

Since we had two runs of sequencing data for the same puck, we were able to compare the effects of the Syrah to the effects of additional sequencing. Using Syrah on the original single sequencing run data resulted in 15.1% more UMIs, 21.3% more beads, and 6.4% more features, while using the standard pipeline with an additional round of sequencing gained only 8.7% UMIs, 3.4% beads, and 3.6% features. [Fig 4B,C]. Syrah provided roughly twice the gains in UMIs and features that additional sequencing did, but resulted in more than ten times the gains in beads, emphasizing the importance of bead de-forking. Using Syrah on the two sequencing run data resulted in similar gains relative to the single run data (13.5% UMIs, 22.9% beads, 6.3% features).

**Figure 4:**
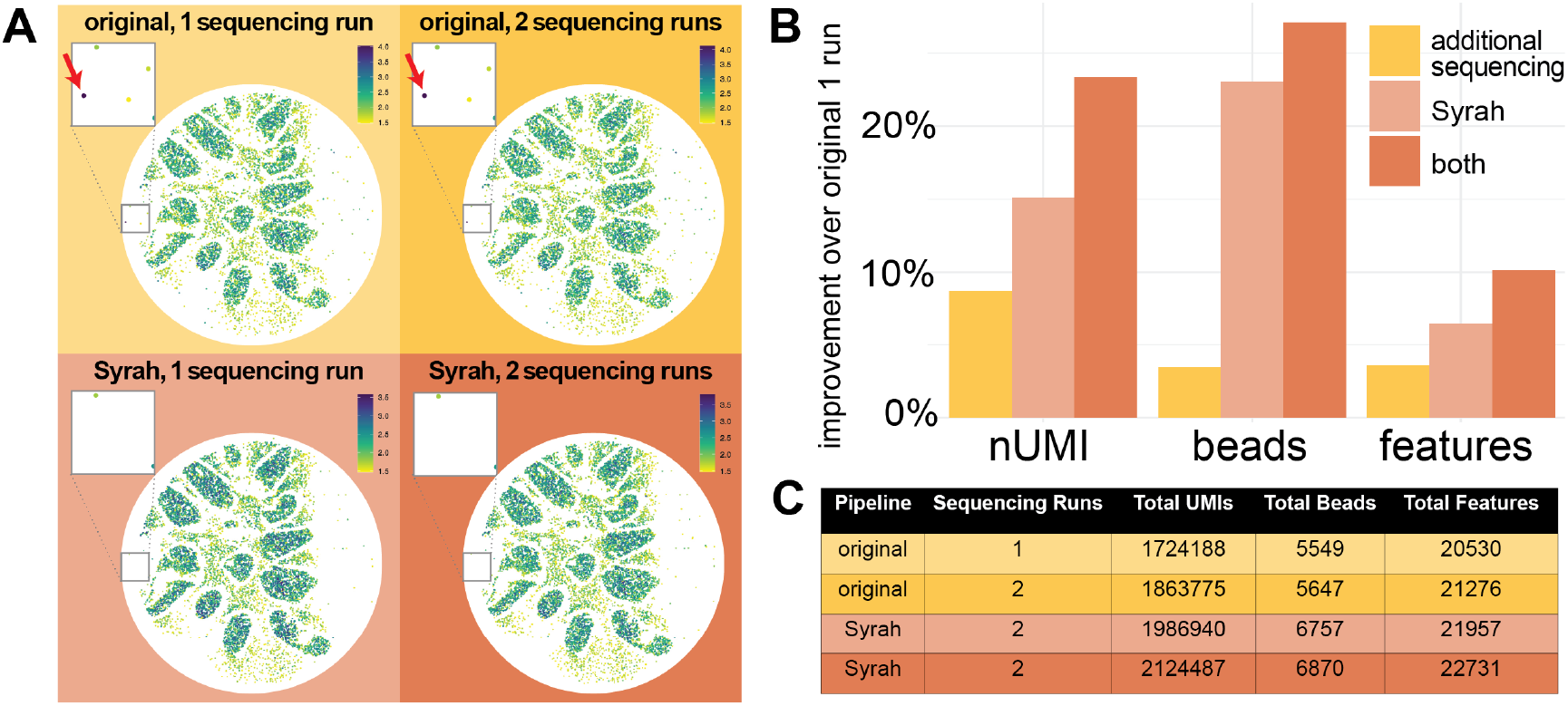
(A) Comparison of results of Syrah and additional sequencing, beads colored by log_10_ nUMI. Red arrows in insets indicate beads with the barcode TTTTTTTTTTTTTT. (B) Percent improvement in UMIs, beads, features over the “original 1 run” reads for additional sequencing, using Syrah, and the combination of both. (C) Summary statistics for all four versions of the data.

On the puck the bead with the barcode TTTTTTTTTTTTTT falls far outside any tissue fragment, but in both non-Syrah datasets it has highest nUMI of any bead, with roughly twice the nUMI of the next highest [Fig 4A top row, inset]. This bead barcode does not appear in either dataset processed with Syrah [Fig 4A bottom row, inset].

## Conclusions

The standard Slide-seqV2 protocol has two sources of errors, oligonucleotide deletions and computational bead “forking”, which reduce both the quantity and quality of the resulting gene expression data.

The improvements provided by Syrah directly correlate with the deletion rate for a given puck, such that pucks with the highest deletion rates and lowest initial quality are most likely to benefit from re-analysis. We hope that this may allow for the salvaging of previously un-interpretable data.

The main drivers of errors addressed here are 1) failed nucleotide incorporation during the synthesis of oligonucleotide sequences used to identify groups of reads, and 2) ambiguities during SOLiD sequencing to determine bead position. Such errors are thus likely to occur with any single-cell or spatial transcriptomic RNA-seq technologies that rely on similar techniques.

## Acknowledgements

We thank Fei Chen and Evan Murray for providing spatially barcoded pucks and experimental support, the Stowers Planaria Core for support with animal husbandry, Anoja Perera for experimental and logistical assistance, the Stowers Sequencing & Discovery Genomics Core for next generation sequencing, the Stowers Computational Biology Core for NGS data processing, and members of the Sánchez Alvarado lab – particularly Enya Dewars – for helpful discussions and feedback.

A.S.A. is an investigator of the Howard Hughes Medical Institute (HHMI) and the Stowers Institute for Medical Research. B.W.B.-P. is a Jane Coffin Childs Memorial Fund Postdoctoral Fellow. F.G.M. is a HHMI Postdoctoral Fellow. This work was supported in part by NIH R37GM057260 to A.S.A.

## SUPPLEMENTARY MATERIALS

### 1. Choice of linker alignment distance cutoff

As linker alignment distance increases, the number of reads where the linker “aligns” steadily decreases until we reach a distance of 6, where it nearly doubles over the distance 5 values [Table S1]. Additionally, the percentage of reads that successfully map to a barcode decreases steadily, but decreases sharply at distance 6 [Table S1]. Taken together these indicated that distance 6 is the point where spurious linker “alignments” overtake actual alignments, so we chose a maximum linker alignment distance of 5.

**Supplementary Table 1:** Read count, barcode part 1 deletion, and barcode mapping information for two-sequencing run data with various linker alignment distances.

| Distance | Mapped | Unmapped | BC1 deletion | Total | % Mapped | % BC1 deletion | % of all reads |
| --- | --- | --- | --- | --- | --- | --- | --- |
| chaotic | 0 | 32501328 | 0 | 32501328 | 0.00 | 0.00 | 17.12 |
| 0 | 105493610 | 27110571 | 26646528 | 132604181 | 79.56 | 20.09 | 69.84 |
| 1 | 6056406 | 3688815 | 2690975 | 9745221 | 62.15 | 27.61 | 5.13 |
| 2 | 794906 | 2559981 | 995604 | 3354887 | 23.69 | 29.68 | 1.77 |
| 3 | 359586 | 1445264 | 633038 | 1804850 | 19.92 | 35.07 | 0.95 |
| 4 | 283068 | 1460837 | 627559 | 1743905 | 16.23 | 35.99 | 0.92 |
| 5 | 263070 | 1372880 | 641368 | 1635950 | 16.08 | 39.20 | 0.86 |
| 6 | 259174 | 2467620 | 936857 | 2726794 | 9.50 | 34.36 | 1.44 |
| 7 | 204734 | 2883110 | 1120572 | 3087844 | 6.63 | 36.29 | 1.63 |
| 8 | 113202 | 557502 | 492088 | 670704 | 16.88 | 73.37 | 0.35 |
| all | 113827756 | 76047908 | 34784589 | 189875664 | 59.95 | 18.32 | 100.00 |

### 2 Proof for deletion rate equation

Let *R_del_* be the deletion rate during oligonucleotide sequencing and *T*_43_ the proportion of thymines at position 43 in read 1. For a given deletion rate *R_del_*, the probability of successful nucleotide incorporation is (1 *− R_del_*) and so the probability of a perfect read, i.e. a read with no deletions through position 43 is (1 *− R_del_*)^43^. Consequently the probability of at least one deletion in positions 1 through 43 is 1 *−* (1 *− R_del_*)^43^. Given that the expected value of *T*_43_ (assuming no deletions) is 25% or 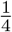, the amount of thymine at position 43 due to perfect reads is 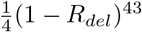 and due to read with 1 or more deletions is 1 *−* (1 *− R_del_*)^43^).

We then see that

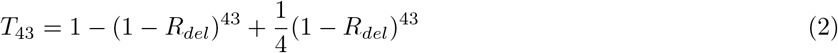

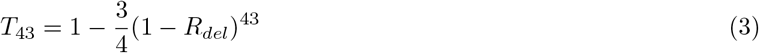

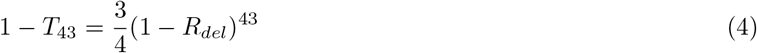

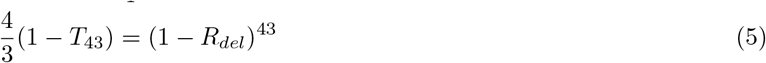

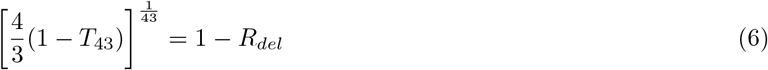

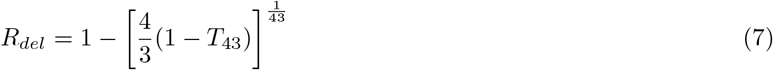

### 3 Notes on *vs1* and *vs2* Slide-seqV2 beads

Slide-seqV2 used two slightly different versions of the bead capture oligo [6], referred to as *vs1* and *vs2*. When specific details differ, such as in Figure 1 and the deletion rate equation, this paper refers to *vs1*. For *vs2* data, the nucleotide at position 41 should be “an A, C or G but not T,” we can estimated the deletion rate of *vs2* data using the proportion of thymines at position 41 (*T*_41_) as

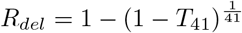

When using Syrah on *vs2* data, simply specify the version as “vs2” in the manifest.

## References

[1] S.L. Beaucage and M.H. Caruthers. “Deoxynucleoside phosphoramidites—A new class of key intermediates for deoxypolynucleotide synthesis”. In: Tetrahedron Letters 22.20 (1981), pp. 1859–1862. issn: 0040-4039. doi: https://doi.org/10.1016/S0040-4039(01)90461-7. url: https://www.sciencedirect.com/science/article/pii/S0040403901904617.

[2] L.J. McBride and M.H. Caruthers. “An investigation of several deoxynucleoside phosphoramidites useful for synthesizing deoxyoligonucleotides”. In: Tetrahedron Letters 24.3 (1983), pp. 245–248. issn: 0040-4039. doi: https://doi.org/10.1016/S0040-4039(00)81376-3. url: https://www.sciencedirect.com/science/article/pii/S0040403900813763.

[3] Kevin Judd McKernan et al. “Sequence and structural variation in a human genome uncovered by short-read, massively parallel ligation sequencing using two-base encoding”. In: Genome Research 19.9 (2009), pp. 1527–1541. doi: 10.1101/gr.091868.109. eprint: http://genome.cshlp.org/content/19/9/1527.full.pdf+html. url: http://genome.cshlp.org/content/19/9/1527.abstract.

[4] Peter W. Reddien and Alejandro Sánchez Alvarado. “FUNDAMENTALS OF PLANARIAN REGENERATION”. In: Annual Review of Cell and Developmental Biology 20.1 (2004). PMID: 15473858, pp. 725–757. doi: 10.1146/annurev.cellbio.20.010403.095114. eprint: https://doi.org/10.1146/annurev.cellbio.20.010403.095114. url: https://doi.org/10.1146/annurev.cellbio.20.010403.095114.

[5] Samuel G. Rodriques et al. “Slide-seq: A scalable technology for measuring genome-wide expression at high spatial resolution”. In: Science 363.6434 (2019), pp. 1463–1467. doi: 10.1126/science.aaw1219. eprint: https://www.science.org/doi/pdf/10.1126/science.aaw1219. url: https://www.science.org/doi/abs/10.1126/science.aaw1219.

[6] Robert R. Stickels et al. “Highly sensitive spatial transcriptomics at near-cellular resolution with Slide-seqV2”. In: Nature Biotechnology 39.3 (2021), pp. 313–319. doi: 10.1038/s41587-020-0739-1. eprint: https://doi.org/10.1038/s41587-020-0739-1. url: https://doi.org/10.1038/s41587-020-0739-1.

[7] Kimberly C Tu et al. “Egr-5 is a post-mitotic regulator of planarian epidermal differentiation”. In: Elife 4 (2015), e10501.

